# Harnessing gene-gene interactions via an RNA-Sequencing network analysis framework improves precision medicine prediction in rheumatoid arthritis

**DOI:** 10.1101/2021.12.08.471133

**Authors:** Elisabetta Sciacca, Anna E. A. Surace, Salvatore Alaimo, Alfredo Pulvirenti, Felice Rivellese, Katriona Goldmann, Alfredo Ferro, Vito Latora, Costantino Pitzalis, Myles J. Lewis

## Abstract

The study of gene-gene interactions in RNA-Sequencing (RNA-Seq) data has traditionally been hard owing the large number of genes detectable by Next-Generation Sequencing (NGS). However, differential gene-gene pairs can inform our understanding of biological processes and yield improved prediction models. Here, we utilised four well curated pathway repositories obtaining 10,537 experimentally evaluated gene-gene interactions. We then extracted specific gene-gene interaction networks in synovial RNA-Seq to characterise histologically-defined pathotypes in early rheumatoid arthritis patients. Specific gene-gene networks were also leveraged to predict response to methotrexate-based disease-modifying anti-rheumatic drug (DMARD) therapy in the Pathobiology of Early Arthritis Cohort (PEAC). We statistically evaluated the differential interactions identified within each network using robust linear regression models, and the ability to predict response was evaluated by receiver operating characteristic (ROC) curve analysis.

The analysis comparing different histological pathotypes showed a coherent molecular signature matching the histological changes and highlighting novel pathotype-specific gene interactions and mechanisms. Analysis of responders vs non-responders revealed higher expression of apoptosis regulating gene-gene interactions in patients with good response to conventional synthetic DMARD. Detailed analysis of interactions between pairs of network-linked genes identified the *SOCS2/STAT2* ratio as predictive of treatment success, improving ROC area under curve (AUC) from 0.62 to 0.78. In conclusions, we demonstrate a novel, powerful method which harnesses gene interaction networks for leveraging biologically relevant gene-gene interactions leading to improved models for predicting treatment response.

## INTRODUCTION

Differential gene expression analysis is a common starting point for many gene expression studies. However, this only reveals differences at the level of individual genes. The identification of interactions between pairs of genes can enhance understanding of biological processes and functional mechanisms which are active within tissues. However, the large number (20,000-50,000) of expressed genes detectable by RNA-Seq renders analysis of all possible gene-gene correlations (of the order of 10^9^) computationally time consuming and confounded by substantial numbers of false positive gene-gene pairs which are biologically and functionally unrelated.

In the current study, we developed a novel network tool integrating information from four pathway repositories(Kanehisa and Goto 2000; Hsu et al. 2011; Xiao et al. 2009; Tong et al. 2019) obtaining 10,537 experimentally evaluated gene-gene interactions. This tool was applied to RNA-Seq data on synovial biopsies from early rheumatoid arthritis (RA) patients to characterise differences in gene-gene interaction networks between histologically defined RA subgroups known as pathotypes(Lewis et al. 2019; Humby et al. 2019), and compare differences between responders and non-responders to conventional synthetic disease-modifying anti-rheumatic drugs (csDMARD) (see Figure 1A).

**Figure 1.**
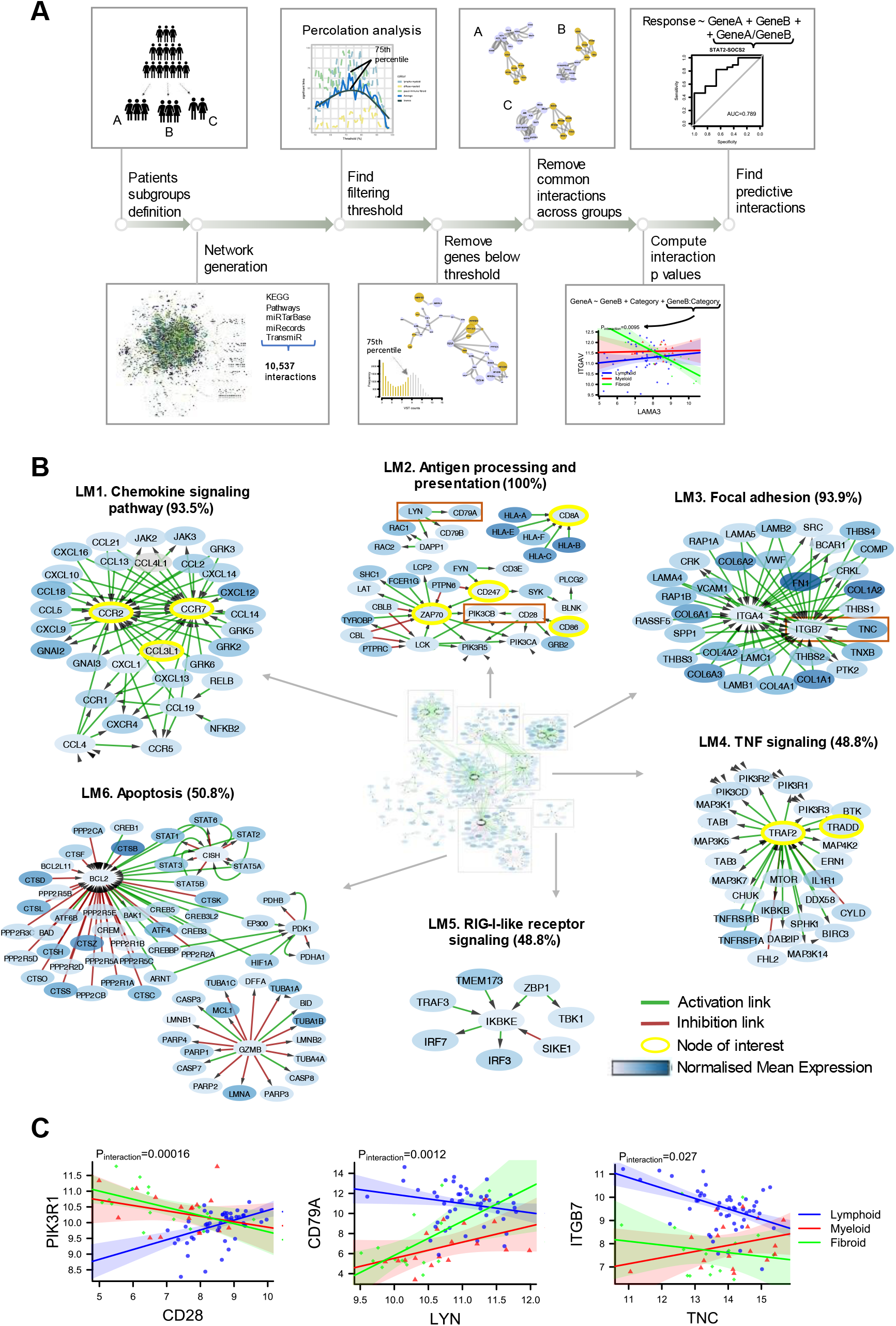
Network analysis of synovial RNA-Sequencing in early RA reveals gene-gene interactions uniquely linked to the lympho-myeloid pathotype. **(A)** Analytical pipeline using network approach to extract informative networks and predictive gene pairs from RNA-seq profiles. Having defined subgroups of patients, an extensive network of interactions is built using merged KEGG pathways enriched with micro-RNAs and transcription factors. The network is replicated for each subgroup and the average expression level of each gene in a subgroup is used to infer a weight on each network node. A first filtering step removes, from each network, nodes (genes) whose weight (subgroup average expression level) is below an optimal threshold obtained via percolation analysis. The second filtering step pull out links (gene-gene interactions) overlapping two or more networks. Robust linear regression with interaction term is used to extract significant gene-gene links. A logistic regression model is built for each significant gene-gene pair to predict response. Ability to predict response is tested by receiver operating characteristic (ROC) curve analysis. **(B)** Network of unique active interactions in the lympho-myeloid pathotype. **Clusters LM1-LM4**. Selected clusters of interest. Labels are determined by gene ontology(GO)/pathway enrichment analysis. Percentages indicate the number of cluster genes included in the associated GO/pathway term. **Cluster LM1**. Cluster of chemokines needed for leukocyte recruitment (93.5% enrichment). **Cluster LM2**. Antigen processing and presentation with T-cell activation genes (100% enrichment). **Cluster LM3**. Group of focal adhesion genes comprising collagens, integrins and laminins (93.9% enrichment). **Cluster LM4**. TNF signaling through mTOR (48.8% enrichment). **Cluster LM5**. Interferon regulation signaling (87.5% enrichment). **Cluster LM6**. Genes of the intrinsic and extrinsic apoptotic pathways (50.8% enrichment). **(C)** Correlation plots showing differential gene-gene correlations with interactions associated with pathotype. Statistical analysis by robust linear regression model. P-value of the gene:pathotype interacting term is shown. Correlation plots of gene pairs CD28 and PIK3R1, CD79A and LYN, and TNC and ITGB7 across different pathotypes.

Patients treated with csDMARD are often subject to lack of treatment efficacy(van der Heijden et al. 2007; Mittal et al. 2012). Several studies have tried to predict patients’ responsiveness based on synovial gene expression, mostly from joint replacement tissue. However, the presence of concomitant immunosuppressive medications and use of microarrays are major limitations of these studies(Badot et al. 2009; Lindberg et al. 2010; De Groof et al. 2016; Mandelin et al. 2018).

In 2019, Lewis et al.(Lewis et al. 2019) used RNA-Seq of synovial biopsies to profile a cohort of patients with early, treatment-naïve RA, and identified three histological and molecular subgroups characterised by i) B-cell infiltration (*lympho-myeloid* pathotype), ii) macrophage infiltration (*diffuse-myeloid* pathotype) and iii) absence of immune cells with stromal cell predominance (*pauci-immune fibroid* pathotype). In the present study we identify functionally relevant gene-gene interactions which are significantly associated with response to csDMARD at six months through robust linear modelling incorporating interaction terms. Significant gene-gene ratios were used to improve predictive models of response to csDMARD treatment as tested by receiver operating characteristic (ROC) curve analysis.

## RESULTS

### Lympho-myeloid pathotype gene network is associated with leukocyte chemokines, chemoattractants and antigen processing

Synovial biopsies from patients with RA demonstrate distinctive histological pathotypes associated with corresponding gene signatures(Lewis et al. 2019; Humby et al. 2019). In the present analysis gene-gene interaction networks specific for the lympho-myeloid, diffuse myeloid and pauci-immune fibroid pathotypes were generated (Figure 1A, see Supplementary Methods for more detail). In brief, an extensive network of curated gene-gene interactions was built by merging KEGG pathways with micro-RNA and transcription factor databases. Average gene expression in each subgroup was used to infer weights on network nodes. Networks were then filtered by percolation analysis based on node weights, removing less highly expressed genes. Interaction pairs overlapping between subgroups were also removed, thus revealing gene-gene networks specific for each subgroup. In Figure 1B the six most prominent clusters of the lympho-myeloid specific network are shown (full network Fig. S1). Gene ontology (GO) enrichment analysis was employed to label gene clusters associated with chemokine signaling (LM1), antigen processing/presentation (LM2), Focal adhesion (LM3), tumor necrosis factor (TNF) signaling (LM4), retinoic acid-inducible gene (RIG)-I-like receptor signaling (LM5), and Apoptosis (LM6) (Table 1).

**Table 1.** GO/Pathway enrichment analysis on network clusters.

| Network | Cluster | Enrichment Term | Nr. of Associated Genes | % of Associated Genes | Adj P-value |
| --- | --- | --- | --- | --- | --- |
| Lympho-myeloid | LM1 | Chemokine signaling pathway | 29/31 | 93.5 | 6.49e-55 |
|  | LM2 | Antigen processing and presentation | 6/6 | 100.0 | 5.78e-14 |
|  | LM3 | Focal adhesion | 31/33 | 93.9 | 2.54e-58 |
|  | LM4 | TNF signaling pathway | 20/41 | 48.8 | 3.72e-32 |
|  | LM5 | RIG-I-like receptor signaling pathway | 7/8 | 87.5 | 1.19e-15 |
|  | LM6 | Apoptosis | 33/65 | 50.8 | 2.74e-52 |
|  | LM7 | MAPK signaling pathway | 44/71 | 62.0 | 1.4e-59 |
|  | LM8 | Wnt signaling pathway | 13/16 | 81.2 | 7.56e-24 |
|  | LM9 | positive regulation of interleukin-8 production | 6/12 | 50.0 | 5.92e-09 |
|  | LM10 | Interleukin-2 family signaling | 3/3 | 100.0 | 2.46e-07 |
|  | LM11 | disulfide oxidoreductase activity | 4/4 | 100.0 | 1.27e-09 |
| Diffuse-myeloid | DM1 | Focal adhesion | 34/45 | 75.6 | 7.1e-57 |
|  | DM2 | PPAR signaling pathway | 22/28 | 78.6 | 1.11e-47 |
|  | DM3 | Dopaminergic synapse | 23/30 | 76.7 | 8.94e-43 |
|  | DM4 | EPHA-mediated growth cone collapse | 3/3 | 100.0 | 3.78e-08 |
|  | DM5 | Cam-PDE 1 activation | 4/4 | 100.0 | 1.41e-13 |
|  | DM6 | Adherens junction | 5/6 | 83.3 | 1.35e-08 |
| Pauci-immune Fibroid | PF1 | Focal adhesion | 42/45 | 93.3 | 2.06e-79 |
|  | PF2 | Ras signaling pathway | 49/64 | 76.6 | 1.09e-79 |
|  | PF3 | Wnt signaling pathway | 23/24 | 95.8 | 1.2e-12 |
|  | PF4 | Cytokine-cytokine receptor interaction | 18/18 | 100.0 | 4.41e-37 |
|  | PF5 | VEGFR2 mediated vascular permeability | 4/7 | 57.1 | 1.07e-06 |
|  | PF6 | Regulation of RUNX1 Expression and Activity | 3/3 | 100.0 | 1.59e-07 |
|  | PF7 | TGF-beta signaling pathway | 9/14 | 64.3 | 6.25e-17 |
|  | PF8 | mTOR signalling | 6/6 | 100.0 | 7.35e-16 |
|  | PF9 | Adrenergic signaling in cardiomyocytes | 9/11 | 81.8 | 5.71e-16 |
|  | PF10 | Vascular smooth muscle contraction | 12/19 | 63.2 | 2.75e-20 |
| Good Responders | R1 | PI3K-Akt signaling pathway | 34/64 | 53.1 | 3.76e-39 |
|  | R2 | ECM-receptor interaction | 7/7 | 100.0 | 2.74e-16 |
|  | R3 | Chemokine receptors bind chemokines | 6/6 | 100.0 | 1.07e-15 |
|  | R4 | MAP2K and MAPK activation | 5/7 | 71.4 | 1.91e-10 |
|  | R5 | disulfide oxidoreductase activity | 4/4 | 100.0 | 1.27e-09 |
|  | R6 | Triglyceride catabolism | 3/3 | 100.0 | 3.77e-08 |
|  | R7 | Cam-PDE 1 activation | 4/4 | 100.0 | 1.41e-13 |
| Non Responders | NR1 | Antigen processing and presentation | 6/6 | 100.0 | 5.78e-14 |
|  | NR2 | Chemokine signaling pathway | 25/25 | 100.0 | 1.6e-49 |
|  | NR3 | Wnt signaling pathway | 23/31 | 74.2 | 1.02e-16 |
|  | NR4 | VEGFR2 mediated vascular permeability | 7/18 | 38.8 | 3.12e-14 |
|  | NR5 | Olfactory transduction | 32/50 | 64.0 | 6.9e-10 |
|  | NR6 | RIG-I-like receptor signaling pathway | 5/6 | 83.3 | 4.88e-31 |
|  | NR7 | Platelet activation, signaling and aggregation | 24/39 | 61.5 | 1.34e-17 |

The presence of multiple chemokine signalling elements including *CCR1* and its ligands *CCL5* and *CCL14, CCR2* together with its ligand *CCL2* and *CCR7* and its ligands *CCL19* and *CCL21*, suggests that local activation and recruitment of lymphocytes to inflamed joints is a central feature of the lympho-myeloid pathotype (Fig. 1B, cluster LM1)(Nerviani and Pitzalis 2018; Firestein and McInnes 2017).

The lympho-myeloid network contained LM2, an antigen processing and presentation cluster (Fig. 1B) with prominence of T-cell genes including *CD8A* with different major histocompatibility complex (MHC) class I genes and T-cell activation genes (*CD247, ZAP70, CD28, CD86*). Using a robust linear model, the lympho-myeloid pathotype showed a significant difference (p=0.00016) in correlation between *CD28* and the class I phosphoinositide 3-kinase (PI3K) signalling regulator *PIK3R1*, which is important for T-cell function downstream of CD28, when compared to the other two pathotypes. B-cell recruitment and stimulation was also implied by the presence of *CXCL13* and the *CD79A-LYN* link (Fig. 1B, cluster LM1). Phosphorylation of the B cell receptor binding CD79a by Lyn kinase is an initial event in B-cell receptor engagement. Differential correlation between *LYN* and *CD79A* was observed in the lympho-myeloid subgroup (p=0.0012) compared to the diffuse-myeloid and pauci-immune fibroid subgroups (Fig. 1C, Table S1).

Invasion and migration of cells requires interaction with the extracellular matrix through macromolecular assemblies known as focal adhesions. Cluster LM3 (Fig. 1B) included multiple focal adhesion genes including collagens, laminins, integrins and Tenascin C (*TNC*), which plays an important role in the development and regeneration of articular cartilage(Hasegawa et al. 2018) and whose interaction with integrins has been widely studied in cancer(Yoshida et al. 2015). Correlation between *ITGB7* and *TNC* was poor in the diffuse-myeloid and pauci-immune fibroid subgroups, but significantly stronger in the lympho-myeloid subgroup (p=0.027).

Other active pathways in the lympho-myeloid pathotype included NF-kB and mammalian target of rapamycin (mTOR) signaling as part of chemokine and TNF signaling (Fig. 1B, clusters LM1 and LM4), and RIG-I-like receptor signaling centred around Inhibitor of nuclear factor kappa B kinase subunit epsilon (*IKBKE)* (cluster LM5). Increased cell turnover in the lympho-myeloid pathotype is suggested by cluster LM6 which contained intrinsic and extrinsic apoptosis related genes *TRAF2, BAD, BAK, BCL2* and cytotoxic T cell marker *GZMB* (granzyme B).

### Macrophage activation and T-cell activation underlie the diffuse-myeloid pathotype gene network

The diffuse-myeloid specific network was of much smaller size (Fig. 2A, full network Fig. S2) after common links were removed, which may reflect the fact that this subgroup has overlapping characteristics with both of the other two pathotypes. On one hand this category is characterized by the infiltration of macrophages, which are also present in the lympho-myeloid subgroup; on the other hand, the absence of B and plasma cell aggregates is a feature in common with the pauci-immune fibroid pathotype.

**Figure 2.**
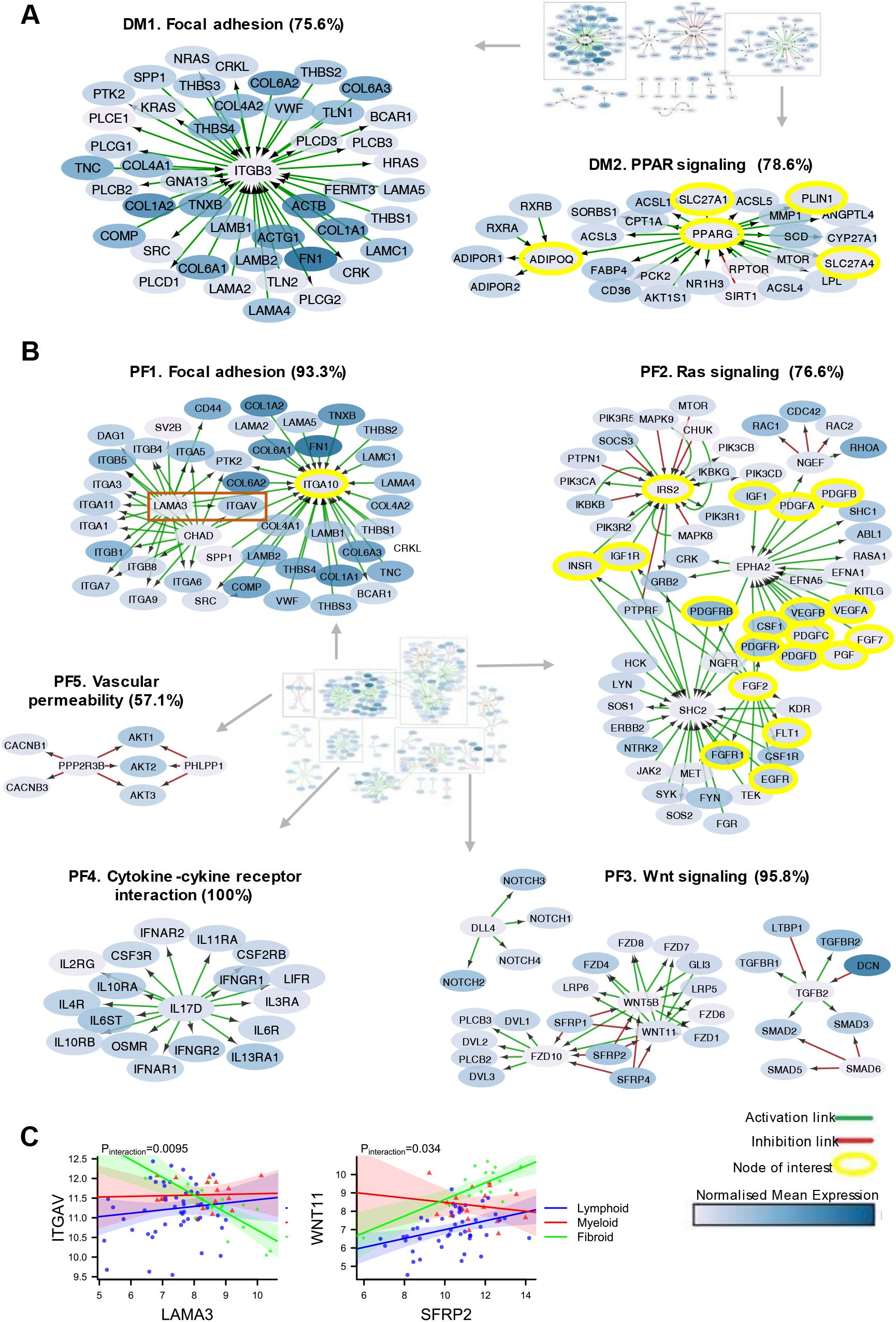
PPAR-γ signaling is key driver of the diffuse-myeloid pathotype while Wnt/Notch signaling pathways characterise the pauci-immune fibroid pathotype. **(A)** Network of unique active interactions in the diffuse-myeloid pathotype. **Cluster DM1**. Extracellular matrix genes for focal adhesion (75.6% enrichment). **Cluster DM2**. Cluster of PPAR signaling pathway (78.6% enrichment). **(B)** Network of unique active interactions in the pauci-immune fibroid pathotype. **Cluster PF1**. Group of focal adhesion genes comprising collagens, integrins and laminins (93.3% enrichment). **Cluster PF2**. Cluster of genes of the Ras signaling pathway (76.6% enrichment). **Cluster PF3**. Clusters of Notch-, Wnt- and TGF-beta signaling (95.8% enrichment). **Cluster PF4**. Cytokine-cytokine interaction of pro-inflammatory genes (100% enrichment). **(C)** Correlation plots showing differential gene-gene correlations with interactions associated with pathotype. Statistical analysis by robust linear regression model. P-value of the gene:pathotype interacting term is shown. Regression plot of ITGAV and LAMA3, WNT11 and SFRP2 in different pathotypes.

One cluster with the same associated GO term as observed in the lympho-myeloid subgroup was for focal adhesion (Fig. 2A, cluster DM1), consistent with the role of integrins in macrophage infiltration into tissues. Uniquely for the diffuse-myeloid pathotype, genes from the PPAR (peroxisome proliferator-activated receptor) signalling pathway, which is involved in fatty acid storage and has been linked to pathological synovial inflammation in RA(Fatel et al. 2018; Arias de la Rosa et al. 2018; Yarwood et al. 2013), were observed in this network (Fig. 2A, cluster DM2). PPAR-γ (*PPARG*), which is critical for macrophage reprogramming(Chawla 2010), and surrounding network genes including adiponectin (*ADIPOQ*) and its receptor (*ADIPOR2*) are significantly upregulated in the diffuse-myeloid pathotype specifically.

### The Wnt signaling pathway characterizes the pauci-immune fibroid pathotype network

Compared to the diffuse-myeloid pathotype, the pauci-immune fibroid pathotype had a more extensive network (Fig. 2B, full network Fig. S3). Extracellular matrix genes including collagens (*COL1A1*, etc.) and laminins (*LAMB1/2*, etc.) were present as an overlapping theme across all three pathotypes in fibroid cluster PF1 (Fig. 2B), lympho-myeloid cluster LM3 (Fig. 1B) and diffuse-myeloid cluster DM1 (Fig. 2A), with different integrins (*ITGA4, ITGB7, ITGB3, ITGA10*) as hubs. Of these, the fibroid hub *ITGA10* is highly expressed by chondrocytes(Bengtsson et al. 2005) and selectively binds collagen(Tulla et al. 2001). Significant statistical interactions across pathotypes were observed for correlations of *ITGA10* and several neighbouring gene nodes (Fig. S4). Another hub node, Chondroadherin (*CHAD*), is a cartilage matrix protein that promotes attachment of chondrocytes, fibroblasts, and osteoblasts(Sommarin et al. 1989). *CHAD* was most strongly correlated with *ITGA10, ITGB4* and *ITGA3* in the lympho-myeloid pathotype (Fig. S5). Among the genes linked to laminin alpha-3 (*LAMA3*), the Integrin Subunit Alpha V (*ITGAV*) is of particular interest since polymorphisms of this gene have been associated with both angiogenesis(Ahnert and Kirsten 2007) and susceptibility to RA(Jacq et al. 2007). In our data its interaction with *LAMA3* showed negative correlation in the pauci-immune fibroid subgroup in contrast to the other two pathotypes (Fig. 2C). These results suggest that *ITGA10, ITGAV, CHAD* and *LAMA3* play central roles in differentiating the pathotypes.

In a separate cluster PF2 we observed several nodes related to the Ras signaling pathway (Fig. 2B), comprising epidermal, fibroblast, nerve, vascular endothelial, insulin-like and platelet-derived growth factors, associated with signaling molecules through Src, *MAPK* and *PI3K*. Fibroblast related pathways were found in cluster PF6 linked to Transforming growth factor (TGF)-beta and Wnt signaling pathway (cluster PF3), which included the secreted frizzled related proteins *SFRP1* and *SFRP2. SFRP1* and *2* are Wnt inhibitors which show reduced expression in RA(Trenkmann et al. 2011; Imai et al. 2006) synovium. Hence it is notable that we found a stronger positive correlation between *SFRP2* and *WNT11* in the fibroid pathotype (Fig. 2C).

Another cluster of major interest was found around the pro-inflammatory cytokine Interleukin (IL)-17D in cytokine-cytokine receptor interaction cluster PF4 (Fig. 2B). IL-17 family members are involved in RA pathogenesis and *IL17D* is expressed in rheumatoid nodules(Stamp et al. 2008). Cluster PF4 comprised cytokine receptors playing key roles in RA, implicating pro-inflammatory activation of fibroblasts and stromal cells in the fibroid pathotype given the absence of immune effector cells. Key transcription factors identified in the fibroid specific network included *RUNX1, AKT, FOXO1A* and mTOR (Table 1). In summary, these analyses revealed functional links between genes characterizing core biological differences which shape each of the three pathotypes.

### Apoptosis genes characterize the good-response network

We performed a separate analysis to identify gene networks related to treatment outcome. Synovial gene expression in patients who responded well to csDMARD was associated with a relatively small gene network (Fig. 3A, full network Fig. S6). The most prominent cluster was centred around B-cell lymphoma 2 (*BCL2*) with genes linked to PI3K-Akt signaling (Fig. 3A, cluster R1). Edges to this node included other cell death regulating genes (*BAX, BAD, BAK1*), multiple cathepsins (*CTSB, CTSS, CTSK*) needed for caspase activation, as well as STAT and mitogen-activated protein (MAP) kinase signaling genes. Additional clusters of genes linked to the good-responder group consisted of alpha and gamma chain laminin genes (cluster R2), and key chemokines and chemokine receptors including *CCL19* and *CXCL13* (cluster R3).

**Figure 3.**
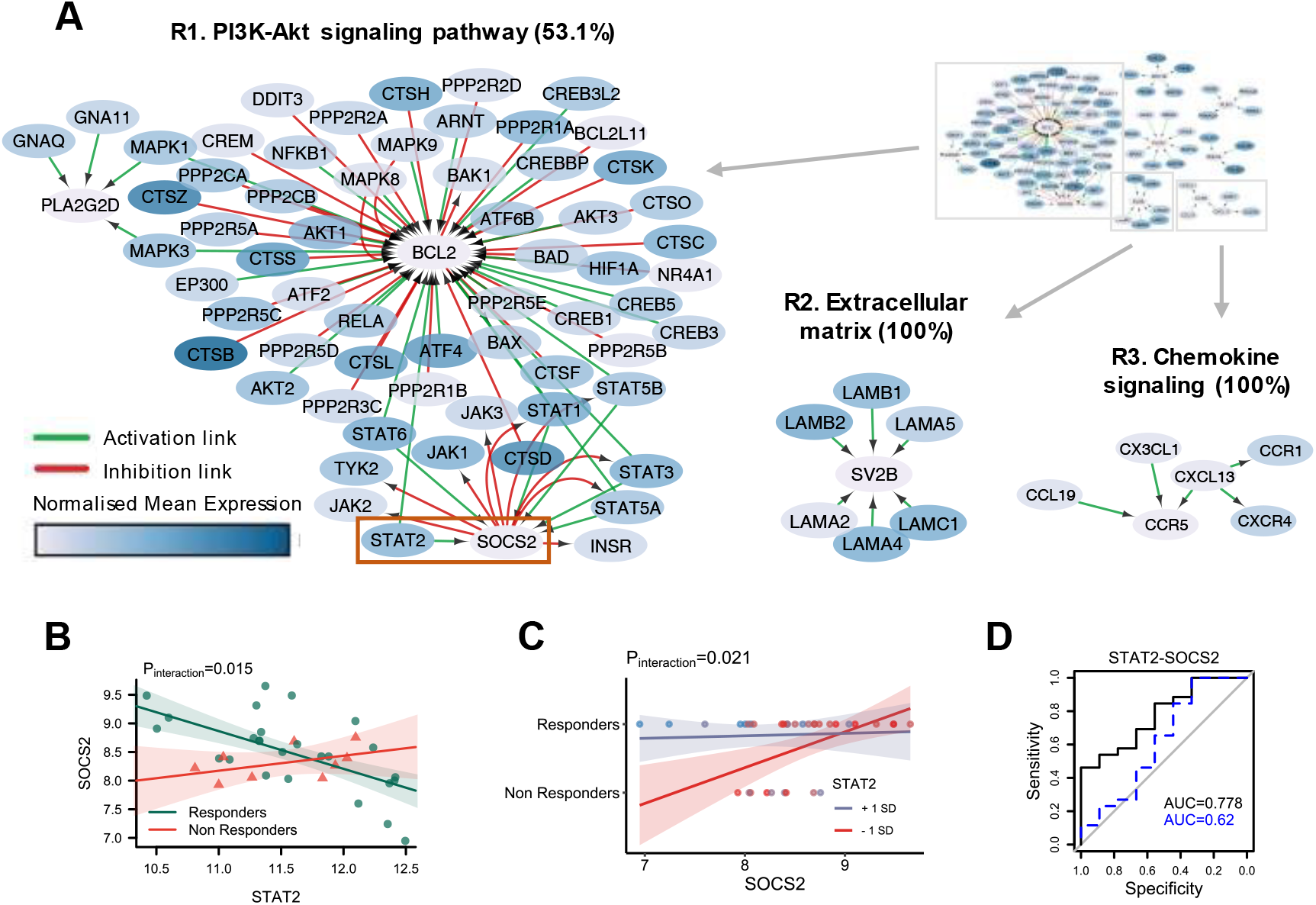
Apoptosis and SOCS/STAT signaling differentiate responders to methotrexate-based therapy from non-responders. **(A)** Network of unique active interactions in conventional synthetic DMARD responders. **Cluster R1**. Cell survival genes part of the PI3K-Akt signaling pathway (53.1% enrichment). **Cluster R2**. Extracellular matrix receptor genes (100% enrichment). **Cluster R3**. Chemokines receptors binding chemokines (100% enrichment). **(B)** Robust linear regression of SOCS2 and STAT2 with interaction term associated with response. P-value of the gene:response interacting term is shown. **(C)** Logistic regression of response as a function of SOCS2 and STAT2. P-value of the response:gene interacting term is shown. Expression of STAT2 is dichotomized at ±1 standard deviation. **(D)** Receiver operating characteristic curve analysis of the response prediction ability of the robust linear model incorporating the ratio term (black line) or not (dotted blue line).

### Activation of the SOCS2-STAT2 negative feedback loop is predictive of good response to csDMARD

Among links characterising the good-responder group, the *SOCS2-STAT2* link is of particular interest. Suppressor Of Cytokine Signaling 2 (*SOCS2*) is one of a family of negative regulators of cytokine receptor signaling that acts on the *JAK/STAT* pathway. Regression analysis revealed differential correlation of *SOCS2* and *STAT2* between responders and non-responders (p=0.015, Fig. 3B, Table S1). Following on from this observation, we fitted a logistic regression model to predict response as a function of the two genes. The interaction term (the gene ratio between *STAT2* and *SOCS2*) was significant (p=0.021) as observed in a plot of the regression model dichotomizing *STAT2* expression at ±1 standard deviation (Fig. 3C). ROC curve analysis of the ability of the model to predict response found that the combined model incorporating *SOCS2, STAT2* and *STAT2*/*SOCS2* ratio showed an area under the curve (AUC) value of 0.78 (Fig. 3D). Removal of the *STAT2*/*SOCS2* ratio term from the linear model resulted in a substantial drop in AUC to 0.62, confirming that the gene ratio interaction term strongly improved the predictive ability of the model.

### Endothelial activation genes link to differential responsiveness to DMARD therapy

The gene expression network specific to the non-responder group showed similarities to the lympho-myeloid gene network as we obtained cluster NR1 of class I human leukocyte antigen (HLA) genes linked by pathway analysis to antigen presentation (Fig. 4A, full network Fig. S7) and cluster NR2 of leucocyte attracting chemokine genes around nodes *CCR2* and *CXCR5*. The B-cell mediator activity of *CXCR5* is known to be initiated by G-protein family genes and particularly depends upon the availability of Gα_i2_ and Gα_i3_. *GNAI3*, which encodes Gα_i3_, showed differential correlation with *CXCR5* when comparing non-responders to responders (Fig. 4B). The non-responder network also showed a Wnt signaling cluster (NR3) analogous to cluster PF3 in the pauci-immune fibroid network, and an angiogenesis cluster, NR4, centred around *NOS3* (Nitric Oxide Synthase 3, eNOS) with surrounding genes linked by pathway analysis to *VEGFR2*-mediated vascular permeability. Vascular endothelial growth factor (VEGF) can activate eNOS either through Ca^2+^/calmodulin or by kinase-mediated phosphorylation(Spiller et al. 2019). We observed a molecular signature for both processes, with differential correlation between *NOS3* and *CAMK1* (calcium/calmodulin dependent protein kinase I) or *AKT3* (AKT serine/threonine kinase 3) with evidence of statistical interaction between responders and non-responders (p=0.036 and p=0.016 respectively, Fig. 4C, D). *AKT1*, a member of the same AKT family that activates *NOS3*, was also found to be correlated with its regulator *PPP2R3B* (protein phosphatase 2 regulatory subunit B’’beta). Statistical analysis by robust linear regression showed a significant interaction term between response and gene expression when comparing response categories (p=0.0096, Fig. 4E). A logistic model incorporating *AKT1, PPP2R3B* and the ratio between the two genes was highly predictive of response (Fig. 4F), reaching an AUC of 0.81 (Fig. 4G). Another Ca^2+^/calmodulin dependent kinase gene, *CAMK2D*, demonstrated differential interaction with *NOS3* between responders and non-responders (p=0.022, Fig. 4H). The *NOS3/CAMK2D* ratio was found to be a significant term (p=0.00904) in a logistic model for prediction of response to csDMARD (Fig. 4I) with an AUC of 0.73, which fell to 0.62 if the *NOS3*/*CAMK2D* ratio term was excluded (Fig. 4J).

**Figure 4.**
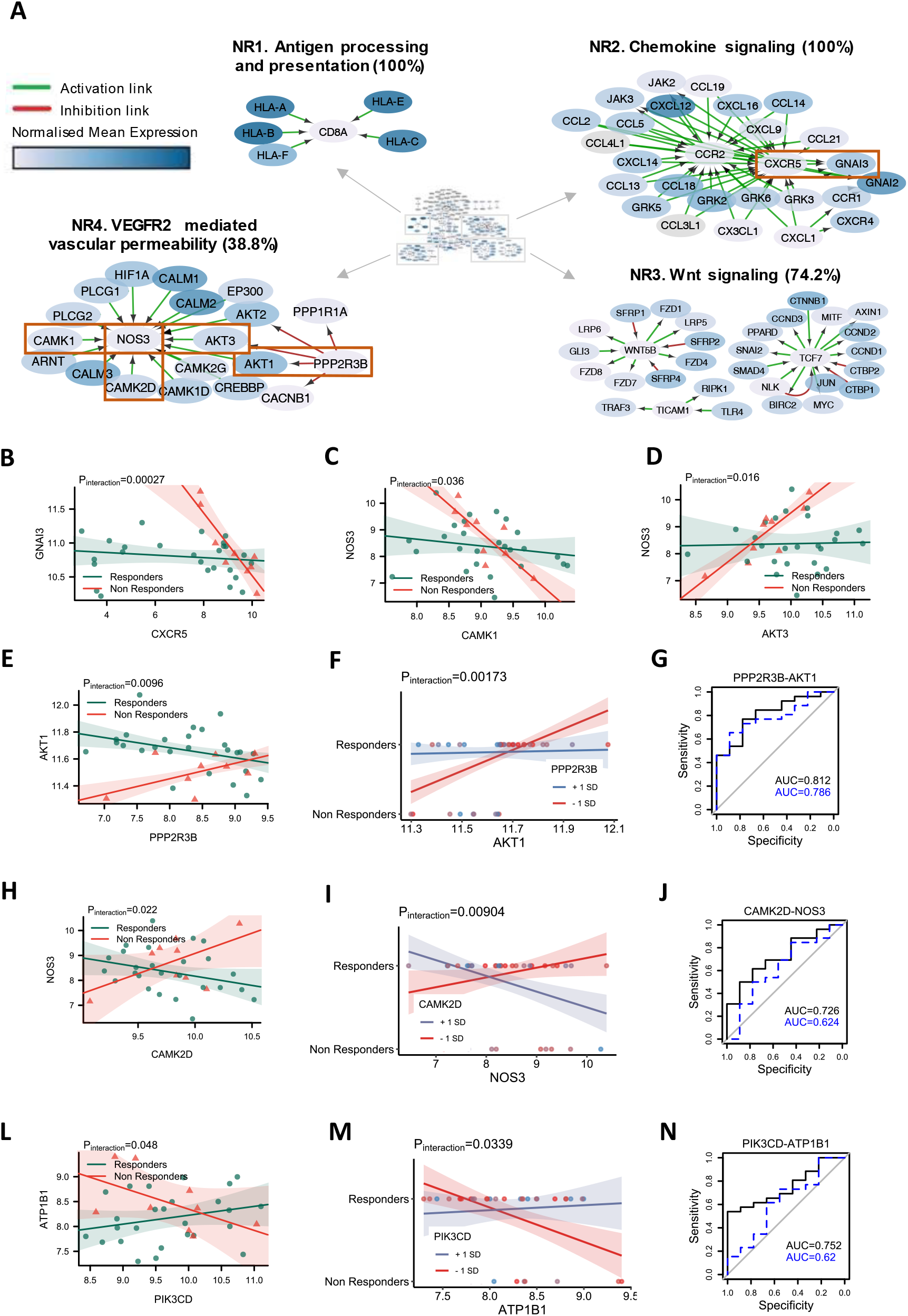
Gene pair interactions linked to endothelial activation and Akt signaling enhance prediction of response to methotrexate-based therapy. **(A)** Network of unique active interactions in conventional synthetic DMARD poor responders. **Cluster NR1**. Antigen processing and presentation cluster (100% enrichment). **Cluster NR2**. Genes of the chemokine signaling pathway (100% enrichment). **Cluster NR3**. Cluster associated to Wnt signaling pathway (74.2% enrichment). **Cluster NR4**. Cluster linked to VEGFR2 mediated vascular permeability (38.8% enrichment). Red boxes highlight predictive gene pairs. **(B-E; H, L)** Robust linear regression with interaction term associated with response for **(B)** GNAI3 and CXCR5 **(C)** NOS3 and CAMK1 **(D)** NOS3 and AKT3 **(E)** AKT1 and PPP2R3B **(H)** NOS3 and CAMK2D **(L)** ATP1B1 and PIK3CD. P-values of the interacting terms are shown. **(F, I, M)** Logistic regression of response as a function of **(F)** AKT1 and PPP2R3B **(I)** NOS3 and CAMK2D **(M)** ATP1B1 and PIK3CD. P-values of the response:gene interacting term are shown. Expression of the second gene is dichotomized at ± 1 standard deviation. **(G, J, N)** Receiver operating characteristic (ROC) curve analysis of the of robust linear model ability to predict response using **(G)** AKT1 and PPP2R3B **(J)** NOS3 and CAMK2D **(N)** ATP1B1 and PIK3CD. All plots show a ROC curve for both the model including the gene-gene ratio interaction term (in black) and the equivalent model excluding the ratio.

Another noteworthy process involved the class I Phosphoinositide 3-kinase (PI3K) gene *PIK3CD* primarily found in leukocytes. Correlation between *PIK3CD* and *ATP1B1* differed significantly between responders and non-responders (p=0.048, Fig. 4L). A logistic regression model fitted to predict response outcome showed a significant interaction term (p=0.0339) for *PIK3CD*/*ATP1B1* ratio (Fig. 4M). The ability of this model in predicting response outcome was good with an AUC of 0.75 (Fig. 4N), which dropped to 0.62 if the interaction term was excluded.

## DISCUSSION

Investigating differential gene-gene interactions in specific groups of patients can enhance understanding of functional pathogenic mechanisms that cannot be captured from single gene level analyses such as differential gene expression studies. Using an extensive network of experimentally validated interactions, we characterised gene networks in histological subgroups (pathotypes) as well as responder/non-responder subgroups of patients from the PEAC cohort(Lewis et al. 2019; Humby et al. 2019). We confirmed our previous finding that the pathotypes are delineated by the type of cells infiltrating the tissue, namely macrophages for the diffuse-myeloid pathotype, B-cells for the lympho-myeloid pathotype and fibroblasts for the pauci-immune pathotype. However beyond this we uncovered critical gene interactions driving processes which clearly differentiate the three pathotypes and may explain underlying mechanisms that differ between pathotypes. Network analysis showed that alterations in the interplay between collagens, laminins and integrins play a central role in differentiating the three pathotypes. This may reflect tissue destructive processes within the joint and extracellular matrix (ECM) remodelling processes leading to differential effects on immune cell tissue infiltration during RA pathogenesis distinguishing the pathotypes.

In lympho-myeloid and the diffuse-myeloid subgroups, the specific gene networks were dominated by TNF and chemokine signalling consistent with macrophage infiltration characterising the diffuse-myeloid subgroup and B/T-cell infiltration characterising the lympho-myeloid subgroup. In addition to these well-known pathways, we observed cytotoxic T-cell genes in the lympho-myeloid network where we observed HLA class I genes around CD8 (Fig. 1B cluster LM2) and pro-inflammatory genes around granzyme B (*GZMB*, cluster LM6), which is a marker for two distinct synovial CD8^+^ T cell subtypes (SC-T5, SC-T6) recently identified in single-cell RNA-Seq studies(Zhang et al. 2019). The diffuse-myeloid gene network (Fig. 2A) demonstrated subnetworks centred around PPAR-γ and its control over fatty acid metabolism which fits with the importance of these pathways in regulating M1/M2 tissue macrophage differentiation. Adiponectin (*ADIPOQ*) has received attention for its role in RA pathogenesis(Fatel et al. 2018) and its expression is elevated in early RA patients(Yoshino et al. 2011; Popa et al. 2009).

In the pauci-immune fibroid pathotype we found gene networks involving i) multiple integrin genes which may represent the interaction between fibroblasts and the ECM, ii) TGF-beta together with SMAD signalling molecules involved in fibroblast differentiation, and iii) an array of growth factor genes (*FGF, PDGF, IGF, VEGF*) and specific cytokines (IL-17D) (Fig. 2B). Thus, the pauci-immune fibroid pathotype consists of an environment driving fibroblast chemotaxis, proliferation and differentiation(Chaudhary et al. 2007).

In the second phase of our analysis we examined networks specific for good-responders to methotrexate-based DMARD therapy in comparison to poor-responders. Multiple chemokines and chemokine receptors were observed in the good-response network, consistent with their importance in immune cell infiltration into inflamed tissues. Humby et al.(Humby et al. 2019) showed that good-responders had significant reduction in synovial expression of genes associated with lymphoid aggregation, as measured by Nanostring panel, including *CXCL13* and *CCL19* which overlap with the present study’s good-response network.

Along with leukocyte recruitment and T-cell activation, the lympho-myeloid subgroup also expressed cluster LM6 which contained apoptosis related genes that showed some degree of overlap with a similar cluster (R1) in the good-responders which was centred around *BCL2*. The role of apoptosis in RA is highly debated(Liu and Pope 2003). One previous study has shown increased caspase activation in inflamed synovial tissue which normalised alongside downmodulation of apoptosis regulators following successful DMARD therapy(Smith et al. 2010).

Multiple STAT and JAK genes were also observed in the good-responder network (Fig. 3A) consistent with their importance in promoting synovial tissue inflammation and the development and mainstream usage of JAK inhibitors as key therapeutics in RA. When analysing gene-gene pairs, we observed a significant interaction between *STAT2* and *SOCS2* expression which differentiated responders and non-responders (Fig. 3B, C). Accordingly, the ratio of *STAT2-SOCS2* significantly improved a prediction model of treatment response (Fig. 3D). A previous study looking at SOCS1-3 found increased expression of *SOCS2* in RA peripheral blood T-cells and synovial fluid macrophages(Isomäki et al. 2007). SOCS genes are typically suppressors of STAT-mediated cytokine signalling, so it is highly plausible that the ratio between specific STAT and SOCS genes could regulate resolution of inflammation and thus influence response to therapy.

This theme of interactions between pairs of genes known to regulate each other was also observed for other gene pairs including *AKT1* and its regulator *PPP2R3B*. Statistical interaction was found between *PPP2R3B* and *AKT1* and response, and the ratio of *PPP2R3B* to *AKT1* improved the AUC of a predictive model. Similarly, *CAMK2D* and *NOS3* showed statistical interaction with response, and the *NOS3/CAMK2D* ratio improved prediction of response. We found similar statistical interactions between *CAMK1* and *NOS3*, and *AKT3* and *NOS3*. These results suggest that altered biological interactions involving these gene pairs differentiates responders from non-responders. The strong involvement of Ca^2+^/calmodulin dependent kinases and eNOS in inflammation-induced vascular permeability suggests that vascular permeability may be a novel mechanism which potentially explains therapeutic response vs failure of methotrexate-based DMARD therapy.

In addition, we also observed notable interactions in other parts of the PI3K/AKT/mTOR pathway, including an improved predictive model for the ratio of PI3 kinase *PIK3CD* and the sodium-potassium ATPase *ATP1B1*. Interestingly, reduction in synovial expression of *PIK3CD* has been linked with response to anti-TNF therapy in RA patients(Marsal et al. 2014; Whitehead et al. 2012). *ATP1B1* has been identified as a biomarker of prognosis and treatment response in different cancer settings(Shi et al. 2016), which suggests that it may be a globally important predictive biomarker.

Our study has limitations i) in the number of patients for which synovial RNA-Seq data was available, and ii) in the lack of similar validation cohorts in early RA. However, we aim to validate the observed gene interactions and predictive models, in future cohorts including the R4RA trial(Humby et al. 2021) and forthcoming STRAP trial (Experimental Medicine & Rheumatology Department 2021).

In summary, we identified gene-gene networks specific to histologically-defined pathotypes in early RA, revealing new biological mechanisms which underlie the development of each pathotype. We identified specific gene networks which differentiate responders to DMARDs from poor-responders. Further analysis of these networks identified gene pairs whose ratios enhanced models predicting response at 6 months. This approach has significant clinical potential to identify interacting pairs of genes which can be used to stratify patients into responders and non-responders.

## METHODS

RNA-Seq data from 94 patients with early, treatment-naïve RA fulfilling the 2010 ACR/EULAR Criteria recruited into the Pathobiology of Early Arthritis Cohort (PEAC)(Lewis et al. 2019; Humby et al. 2019) was used for the current study. 11 samples had ungraded histopathology or were removed due to poor RNA quality, leaving 83 samples in the present study (Table 2). Patients were stratified into three synovial histopathological subgroups: lympho-myeloid, diffuse-myeloid and pauci-immune as previously described(Lewis et al. 2019). After a baseline biopsy (treatment-naïve), patients underwent six months of methotrexate-based csDMARD therapy. Responsiveness was assessed according to DAS28 EULAR criteria. Our study compared both histopathological and treatment response groups, with separate analyses run for each classification (Table 1). The analytical pipeline summarised in Figure 1A shows the steps through which informative gene networks and predictive gene pairs were extracted for each classification (see supplementary methods for full details).

**Table 2.** Baseline demographics of treatment-naïve RA patients recruited into the Pathobiology of Early Arthritis Cohort (PEAC) used for this study.

|  | Lymphoid (N=49) | Myeloid (N=18) | Fibroid (N=16) | Total (N=83) | p value |
| --- | --- | --- | --- | --- | --- |
| **Age (years)** | 52.3 (16.2) | 50.4 (16.5) | 53.2 (15.0) | 52.1 (15.9) | 0.865 |
| **Gender** |  |  |  |  | 0.608 |
| F | 37 (75.5%) | 12 (66.7%) | 13 (81.2%) | 62 (74.7%) |  |
| M | 12 (24.5%) | 6 (33.3%) | 3 (18.8%) | 21 (25.3%) |  |
| **Disease duration (months)** | 5.9 (3.3) | 4.8 (2.6) | 7.0 (3.5) | 5.8 (3.3) | 0.152 |
| **RF** |  |  |  |  | 0.203 |
| pos | 32 (69.6%) | 9 (52.9%) | 7 (46.7%) | 48 (61.5%) |  |
| neg | 14 (30.4%) | 8 (47.1%) | 8 (53.3%) | 30 (38.5%) |  |
| **CCP** |  |  |  |  | 0.025 |
| pos | 39 (84.8%) | 10 (55.6%) | 9 (60.0%) | 58 (73.4%) |  |
| neg | 7 (15.2%) | 8 (44.4%) | 6 (40.0%) | 21 (26.6%) |  |
| **DAS28** | 6.2 (1.2) | 5.6 (1.2) | 5.2 (1.6) | 5.8 (1.3) | 0.029 |
| **ESR (mm/hr)** | 50.9 (28.1) | 37.6 (25.4) | 30.8 (27.7) | 44.1 (28.4) | 0.025 |
| **CRP (µg/mL)** | 25.0 (26.5) | 17.5 (26.0) | 14.4 (41.7) | 21.3 (29.9) | 0.395 |
| **TJC** | 12.6 (7.2) | 9.9 (6.3) | 10.9 (8.9) | 11.7 (7.4) | 0.396 |
| **SJC** | 8.4 (5.7) | 7.1 (4.3) | 5.2 (5.0) | 7.5 (5.4) | 0.116 |
| **VAS** | 67.9 (24.0) | 61.1 (21.7) | 57.8 (27.1) | 64.5 (24.2) | 0.287 |
| **HAQ** | 1.6 (0.8) | 1.4 (0.6) | 1.6 (0.8) | 1.5 (0.7) | 0.499 |
| **DAS28 EULAR** |  |  |  |  | 0.603 |
| Good | 15 (36.6%) | 4 (30.8%) | 7 (53.8%) | 26 (38.8%) |  |
| Moderate | 20 (48.8%) | 8 (61.5%) | 4 (30.8%) | 32 (47.8%) |  |
| None | 6 (14.6%) | 1 (7.7%) | 2 (15.4%) | 9 (13.4%) |  |

## DATA ACCESS

The dataset supporting the conclusions of this article is available in the ArrayExpress repository, https://www.ebi.ac.uk/arrayexpress/experiments/E-MTAB-6141/. The code used for analysis is publicly available on the online github hosting service at https://github.com/elisabettasciacca/DEGGs The code is currently being developed into an R package and will be submitted to the Bioconductor repository shortly.

## ACKNOWLEDGEMENTS

The Pathobiology of Early Arthritis Cohort (PEAC) was supported by funding from the UK Medical Research Council (MRC) (grant number G0800648).

