## Supplemental Methods and Figures for "Harnessing gene-gene interactions via an RNA-Sequencing network analysis framework improves precision medicine prediction in rheumatoid arthritis"

**SUPPLEMENTARY MATERIAL**

### Supplementary Methods

*Patient cohort*

In total, RNA-Seq data from 94 patients with early, treatment-naïve RA fulfilling the 2010 ACR/EULAR Criteria who had been recruited into the Pathobiology of Early Arthritis Cohort (PEAC)(Lewis et al. 2019; Humby et al. 2019) was used for the current study. 11 samples were removed due to poor RNA quality (n=7) or ungraded histopathology (n=4), thus 83 samples were included in the present study (see Table 1). As previously described(Lewis et al. 2019; Humby et al. 2019), patients were stratified into three distinct synovial histopathological groups: (1) *lympho-myeloid* characterized by lymphoid cell infiltration such as T-cells, B-cells and plasma cells together with myeloid cells; (2) *diffuse-myeloid* with a high prevalence of cells from the myeloid lineage with low numbers in B-cells and plasma-cells; (3) *pauci-immune* identified by lack of infiltrating immune cells associated with stromal cell expansion. After a baseline biopsy (treatment-naïve), patients underwent six months of csDMARD therapy and responsiveness was assessed according to DAS28 EULAR criteria (see Table 2). Clinical outcome measures were determined by clinicians entirely blinded to any synovial biopsy histology and RNA-Seq data. All RNA extraction and RNA-Seq was performed independently of knowledge of clinical outcome.

*Network analysis*

The study pipeline is summarized in Figure 1A. Samples from the PEAC study were initially categorised into histopathological and treatment response groups with separate analyses run for each type of classification. After Variance Stabilizing Transformation (VST) of the count data using DESeq2, the mean normalised gene expression profile was derived for each group. The differential gene-gene correlation analysis was restricted to gene pairs whose interactions have been previously experimentally confirmed. Alaimo et al.(Alaimo et al. 2016) previously merged information from four well-curated pathway repositories to generate a network of 10,537 interactions obtained from KEGG(Kanehisa and Goto 2000), mirTARbase(Hsu et al. 2011), miRecords(Xiao et al. 2009) and transmiR(Tong et al. 2019). This network was exported via the *exportgraph* function in MITHrIL(Alaimo et al. 2016) and replicated for each group. To generate networks specific for each histopathological or response subgroup, weights were assigned to gene nodes, equal to the computed mean gene expression across that particular subgroup. Nodes representing genes whose mean expression was below the 75th percentile of the entire VST matrix were removed from each network. This cut-off level was determined following a percolation analysis. Commonly used in statistical physics and mathematics, percolation describes the behaviour of network properties when a certain percentage of nodes or links are removed(Sahini and Sahimi 1994). In this implementation, the percolation threshold was optimised to maximise the number of statistically significant differential interactions.

In a second filtering step remaining interactions in common between two or more group networks were removed, to obtain group-specific interactions for each network.

The resulting set of networks presented easily identifiable clusters that were enriched by means of ClueGO(Bindea et al. 2009) (v.2.5.5). Four repositories were selected for enrichment: KEGG(Kanehisa and Goto 2000), REACTOME(Jassal et al. 2020) and two Gene Ontology (GO) databases(Camon et al. 2004), *BiologicalProcess-EBI-UniProt-GOA* and *ImmuneSystemProcess-EBI-UniProt-GOA* at 5 or 6 GO tree levels.

For each cluster, the significant GO term/pathway ($p < 0.05$) containing the highest percentage of cluster genes was annotated in Table 2. Enrichment percentage scores refer to the percentage of genes within a specific cluster which are enriched within a given pathway.

The code used to implement the described network analysis is publicly available on github (https://github.com/elisabettasciacca/DEGGs) and will be submitted as an R package to the Bioconductor repository shortly.

*Robust linear regression interaction analysis*

We then evaluated the statistical significance of single links by estimating the expression of each node interaction of a group network. A robust linear regression model with interaction term was fitted using the rlm function from the MASS (v.7.3) R package:

$$GeneA_i = \beta_0 + \beta_1 \, GeneB_i + \beta_2 \, Response_i + \beta_3 \, GeneB_i * Response_i + \varepsilon_i$$

where i = 1, …, n, is the number of samples and $\varepsilon_i$ are random variables.

*Response* is replaced by *Pathotype* in the case of pathotype comparison.

Interaction analysis was performed across all gene nodes present within each network (Table S1).

The ratio of gene pairs whose interaction term *GeneB * Response* were significant was used to predict response (as categorical variable) to csDMARD treatment:

$$Response_i = \beta_0 + \beta_1 \, GeneA_i + \beta_2 \, GeneB_i + \beta_3 \, GeneA_i \, / \, GeneB_i + \varepsilon_i$$

Models were tested to determine whether the presence of the interaction term expressed by the ratio of the two genes enhance the prediction ability of the model. Fitted models were tested for predictive ability by likelihood ratio test P-value as well as receiver operating characteristic (ROC) curve analysis using the pROC package in R in comparison to the equivalent model formula without the additional gene/gene ratio term.

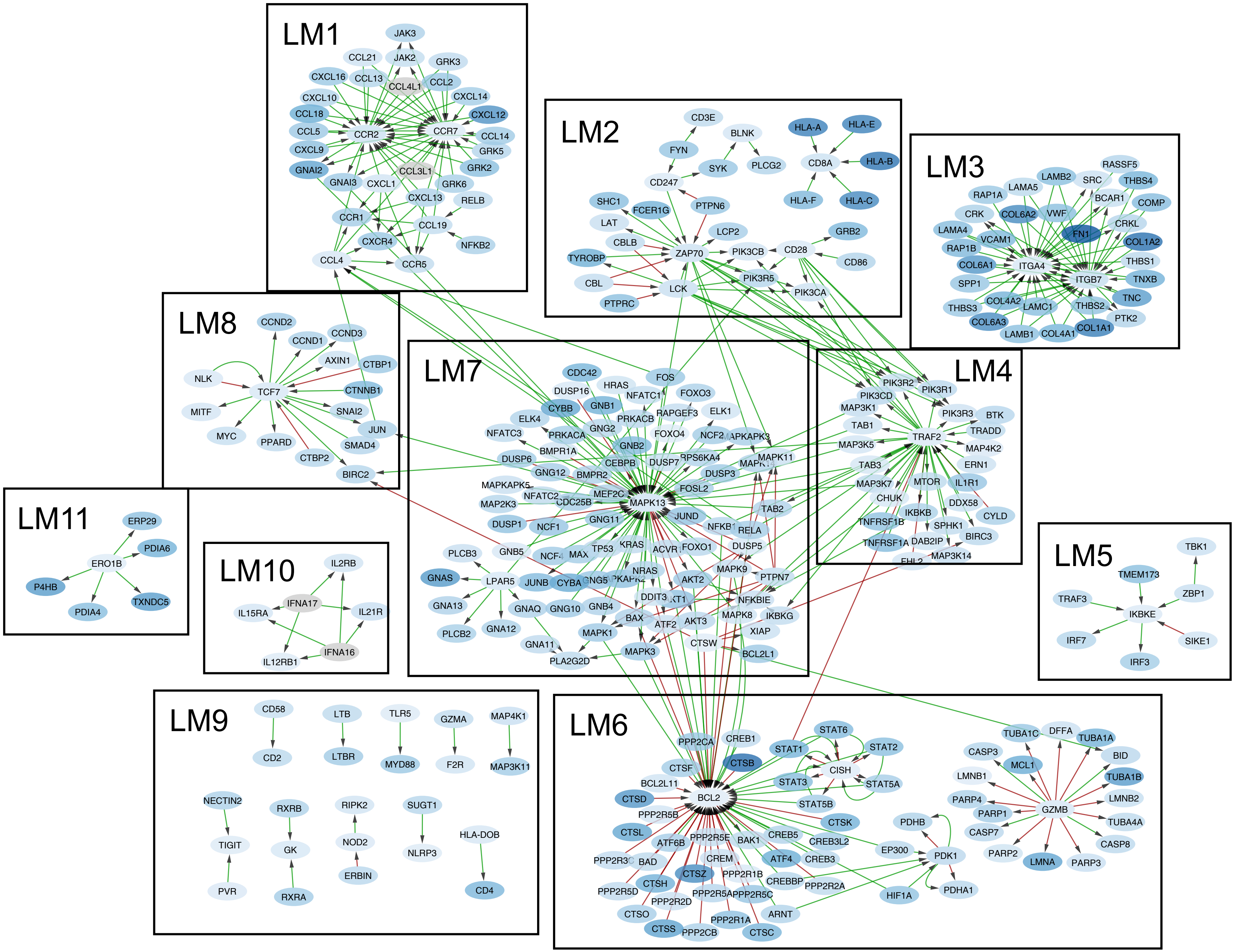


**Figure S1.** Network of unique interactions for the lympho-myeloid pathotype.

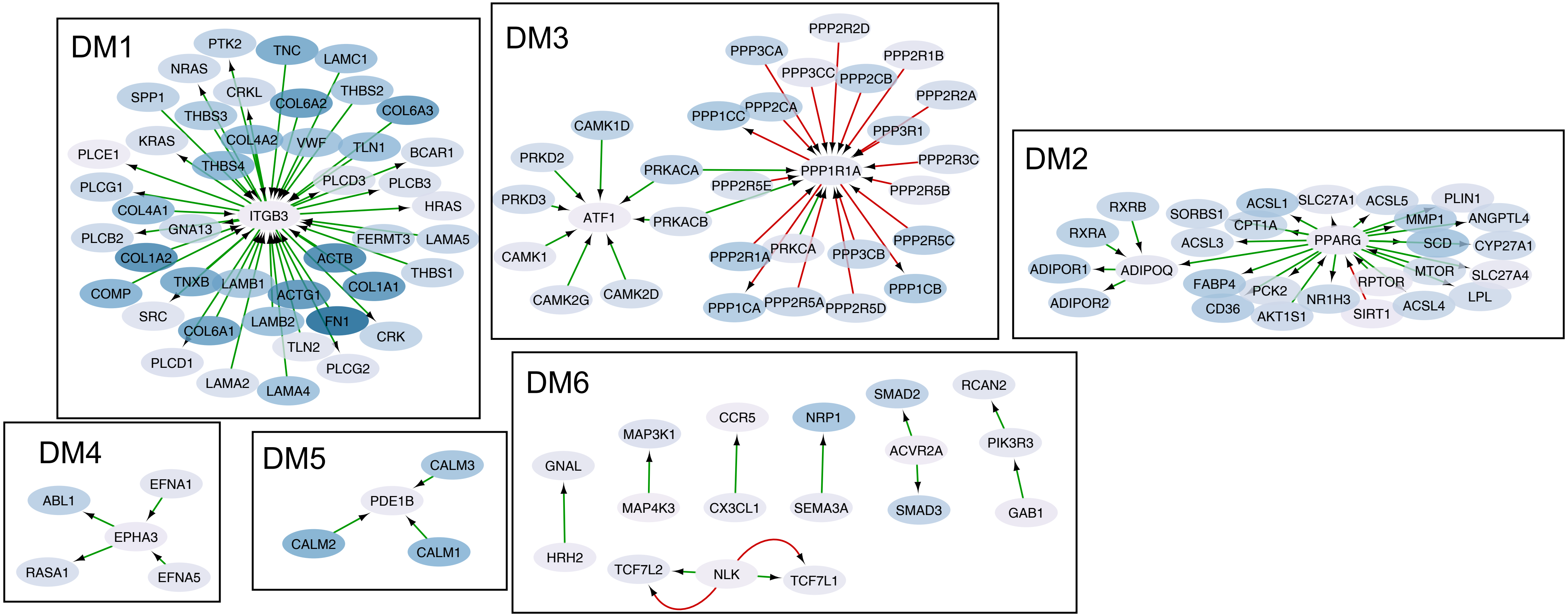


**Figure S2**. Network of unique interactions for the diffuse-myeloid pathotype.

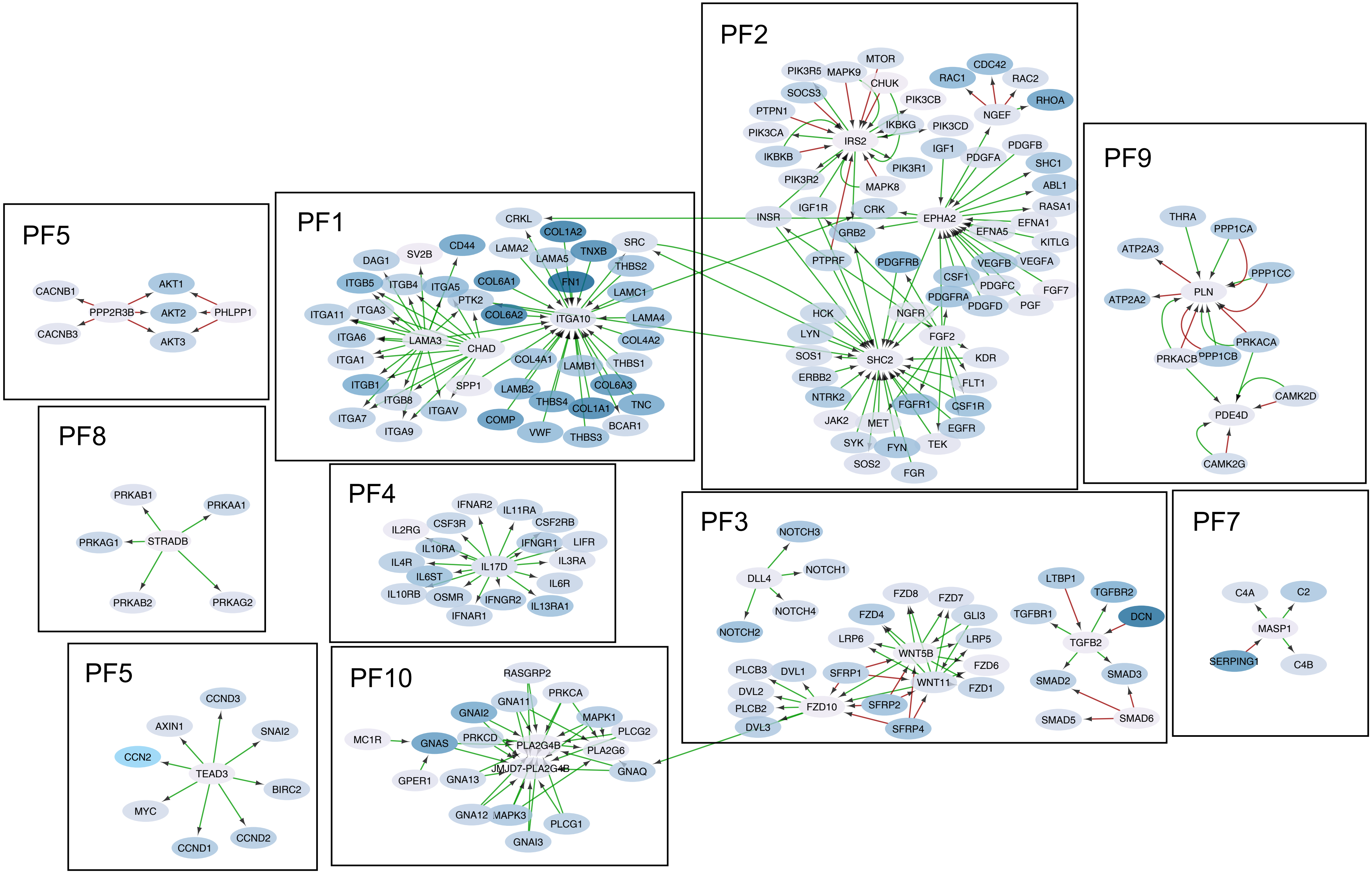


**Figure S3.** Network of unique interactions for the pauci-immune fibroid pathotype.

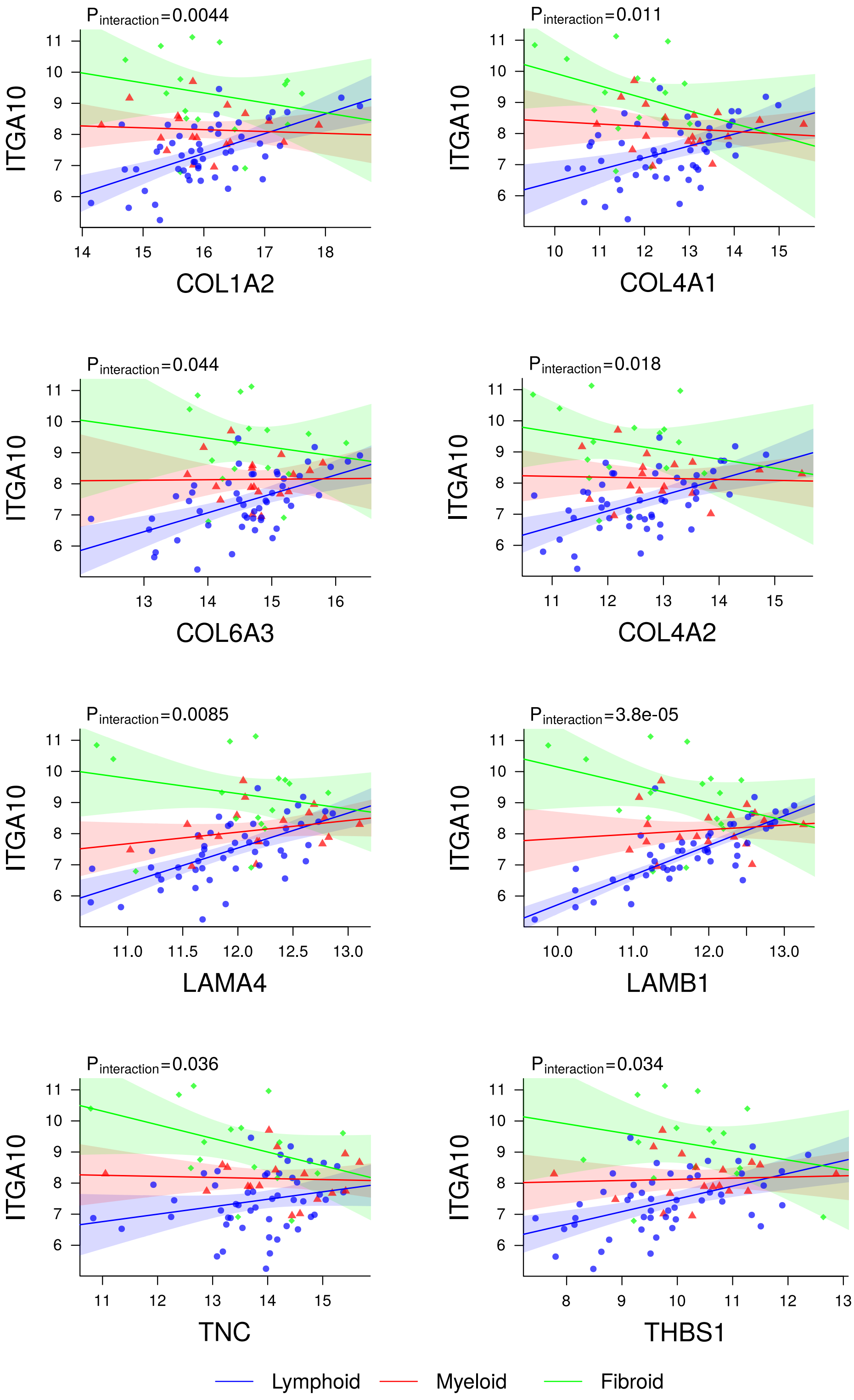


**Figure S4**. Interaction of integrin alpha 10 (*ITGA10*) with ECM genes differentiates the three pathotypes with the same recurring pattern of correlations.

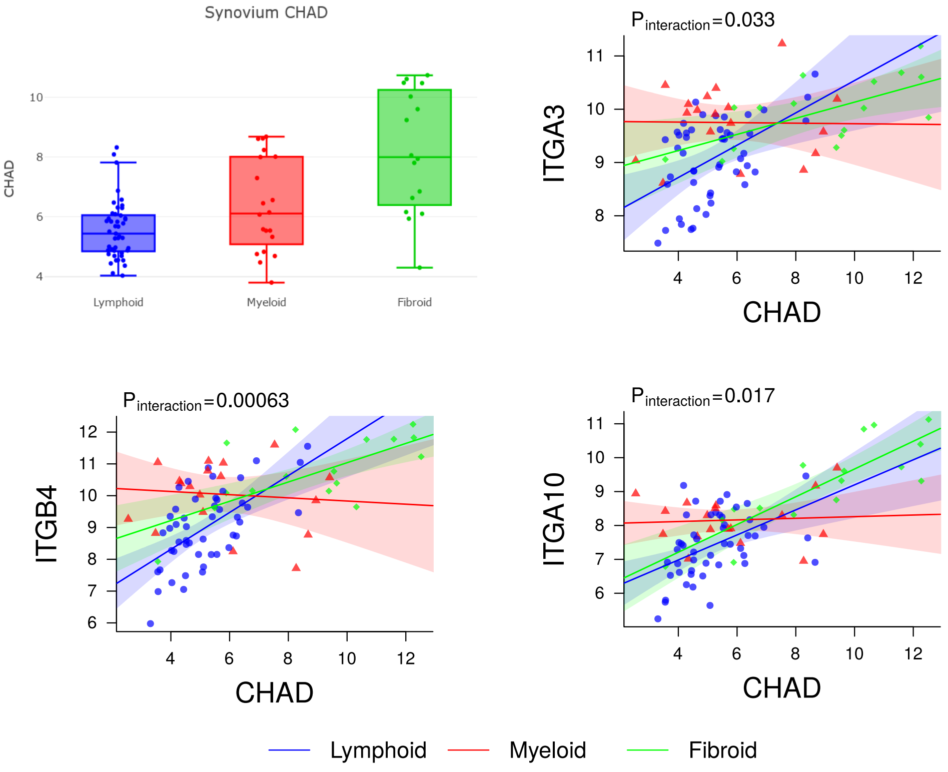


**Figure S5.** Chondroadherin (*CHAD*) shows different levels of synovial expression across pathotypes and its interaction with multiple integrins reflects differential correlations between the histological subgroups.

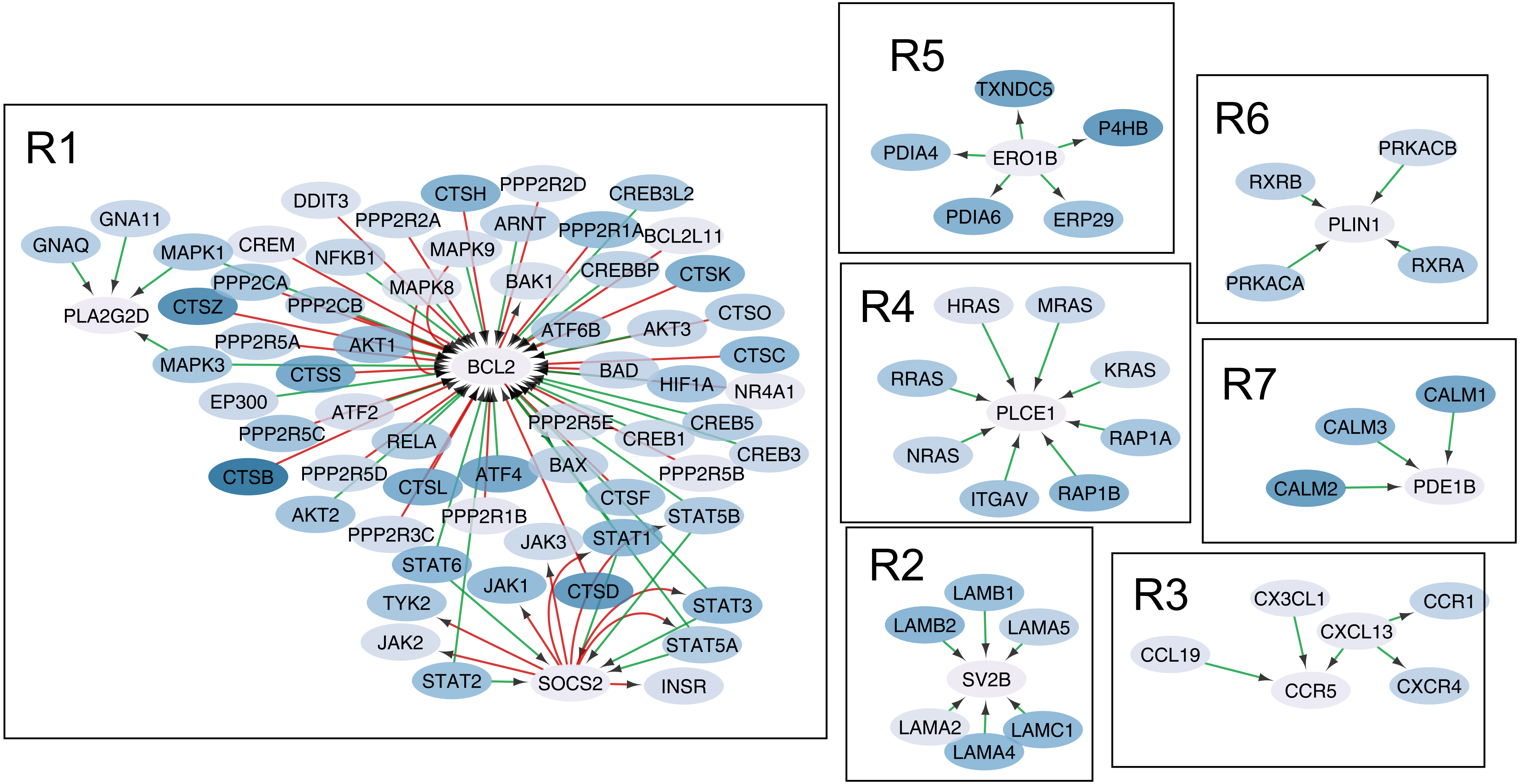


**Figure S6.** Network of unique interactions for the good responders group.

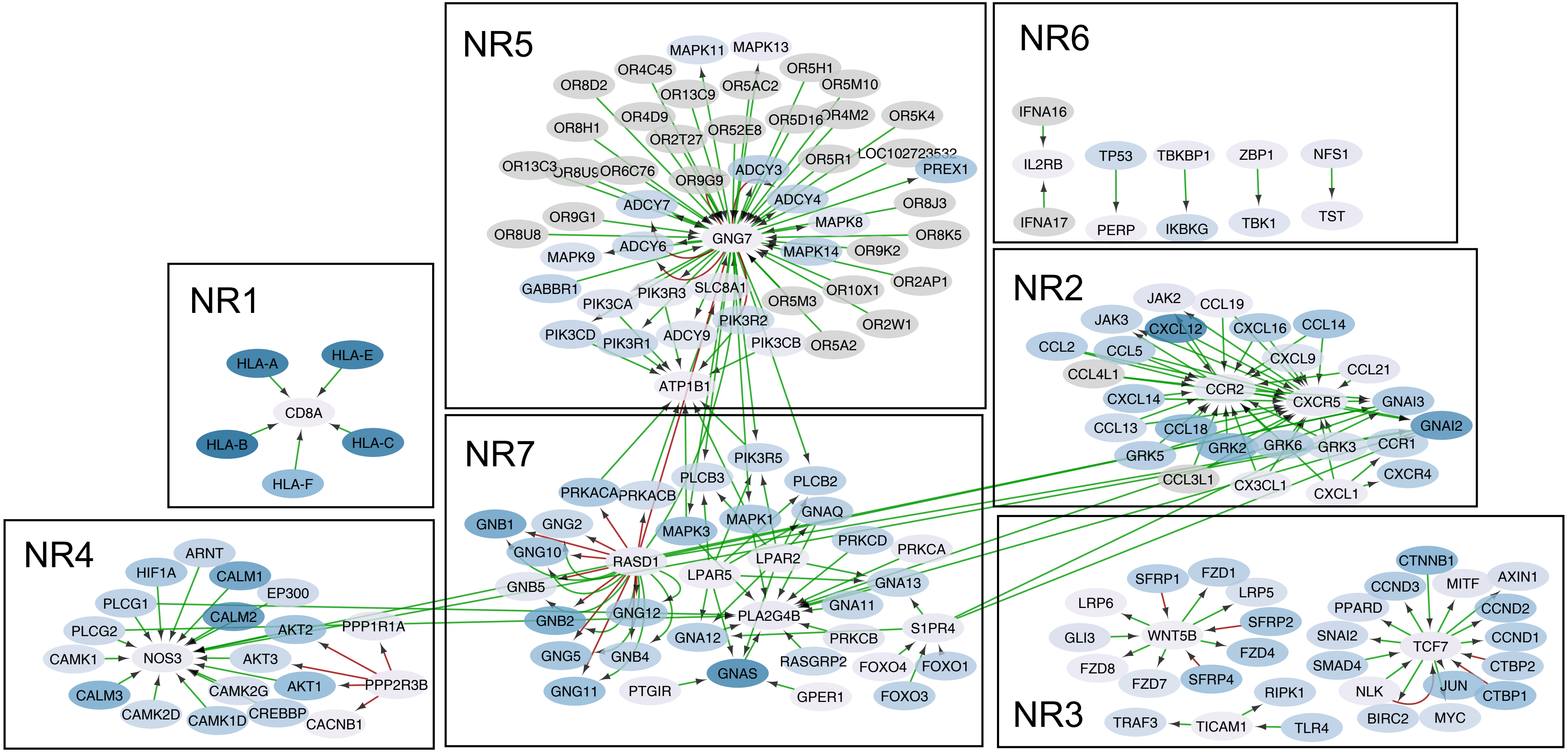


**Figure S7.** Network of unique interactions for the non responders group.
